# Type I Natural Killer T Cells Suppress Infection-Induced Cytokine Storm Through an IL-22-STAT3 Axis

**DOI:** 10.64898/2026.06.14.731971

**Authors:** Cameron M. Torres, Nicole R. Setzu, Brianna Rodriguez, Aylin Sanchez Guillen, Leslie Rodriguez, Noel Molina-Limon, Angel Gutierrez, Luisa Rodriguez, Crystal Devora, Michelle Sanchez Guillen, Charles T. Spencer

## Abstract

Severe infections can trigger systemic inflammatory response syndrome (SIRS), wherein excessive cytokine release generates a “cytokine storm” causing tissue damage, multi-organ failure, and death. Natural killer T (NKT) cells are innate-like lymphocytes that respond rapidly to infection and can either amplify or suppress inflammation. Distinct NKT subsets may have opposing roles in acute infection, but their specific contributions to hyperinflammation remain unclear. Using an intradermal murine *Francisella tularensis* LVS model of infection-induced cytokine storm, we demonstrate that type I NKT cells act as dominant suppressors of systemic hyperinflammation, whereas type II-enriched NKT cells confer limited protection.

This immunoregulation occurs via secreted mediators rather than direct cytotoxicity or cell-cell contact. Notably, we identify IL-22 as a key type I NKT cell effector that suppresses pro-inflammatory cytokine levels. These findings define a novel IL-22-dependent type I NKT cell immunoregulatory axis that restrains pathological macrophage cytokine responses during severe bacterial infection.

## Introduction

Systemic Inflammatory Response Syndrome (SIRS) results from severe bodily stress causing dysregulated inflammation. In response to certain severe infectious diseases (e.g., tularemia, influenza, SARS-CoV-2, Ebola, etc. as well as bacterial sepsis),[1], [2], [3], [4], [5], [6], [7], [8], [9], [10], [11] this dysregulated inflammation results from excessive cytokine production leading to tissue damage, multiorgan dysfunction syndrome (MODS), and death.[12], [13], [14], [15], [16], [17], [18], [19], [20], [21] While inflammatory cytokines are essential for antimicrobial defense, rapid, massive release of these cytokines into the blood, commonly termed a ‘cytokine storm,’ results in Cytokine Release Syndrome (CRS), a form of SIRS.[21], [22], [23], [24], [25], [26] For example, in tularemia, disease severity strongly correlates with elevated levels of pro-inflammatory (IL-6, TNF-α, IFN-γ, IL-17) and regulatory cytokines (IL-10, IL-25, TGF-β) highlighting immune-mediated pathology as a critical determinant rather than bacterial burden alone.[1], [2], [27]

Natural killer T (NKT) cells are a distinct subset of innate-like T lymphocytes that rapidly respond to infection and bridge innate and adaptive immunity.[28], [29] Unlike conventional T cells, NKT cells recognize lipid antigens presented by the non-polymorphic molecule CD1d and can be activated within hours of pathogen exposure.[30], [31], [32], [33], [34], [35] Because of this rapid activation and the diversity of their response, eliciting crosstalk with macrophages, dendritic cells, NK cells, and conventional T cells, they are able to act as early regulators of immune responses.[36], [37], [38], [39], [40] Accumulating evidence indicates that NKT cells play context-dependent roles in inflammation and infectious disease.[39], [40], [41], [42], [43], [44], [45] However, numerous prior studies have examined NKT cells as a single population, without resolving the distinct contributions of type I and type II subsets.

NKT cells are broadly divided into two major subsets.[35], [46], [47], [48] Type I (invariant) NKT cells express a semi-invariant T cell receptor that recognizes α-galactosylceramide and structurally related glycolipids,[30], [49] whereas type II NKT cells express more diverse TCRs and respond to a broader range of endogenous and microbial lipids.[46], [50], [51], [52], [53], [54], [55] In several viral and intracellular bacterial infections, type I NKT cells enhance host defense and modulate myeloid cell function. Indeed, excessive iNKT cell activation has been linked to immunopathology in respiratory infections and sepsis-like syndromes.[56], [57], [58], [59], [60], [61], [62] Type II NKT cells have been implicated in both immune regulation and tissue injury, depending on disease context and tissue localization.[53], [63], [64] These divergent observations suggest that NKT cell function is not uniform but instead shaped by subtype, tissue environment, and antigenic stimulus.[41], [43], [65]

*Francisella tularensis*, the causative agent of tularemia, is a highly virulent intracellular pathogen transmissible by numerous infection routes including aerosol and arthropod exposure.[66], [67], [68], [69] As a model of infection-driven cytokine storm, it is widely used for studying the development and control of the cytokine storm.[1], [2], [27], [67] NKT cells are activated early (within 12-24 hours) during *F. tularensis* infection, but prior studies have yielded conflicting conclusions regarding whether these NKT responses are protective or pathogenic.[42], [59], [69], [70]

Herein, we investigated the subtype-specific roles of type I and type II NKT cells in the *F. tularensis*-induced hyperinflammatory cytokine storm. We demonstrate that in response to intradermal inoculation with *F. tularensis*, type I NKT cells act as dominant suppressors of the cytokine storm both *in vivo* and *in vitro*. This immunoregulation acts through a mechanism mediated by soluble factors rather than cytotoxicity or direct cell–cell contact. We further identify IL-22 as a key effector molecule of type I NKT cell–mediated suppression. These findings define a previously underappreciated immunoregulatory axis and provide mechanistic insight into how NKT cells restrain pathological inflammation during severe bacterial infection.

## Materials and Methods

### Ethics Statement

All animal procedures were carried out in accordance with the Guide for the Care and Use of Laboratory Animals and approved by the Institutional Animal Care and Use Committee (IACUC) under protocol 202407-1.

### Animals

C57BL/6J mice (strain #000664) and B6.129S6-CD1d^-/-^ mice (strain #008881) were obtained from Jackson Laboratories. B6.129-Jα18^-/-^ mice were used with permission of Masaru Taniguchi under MTA with RIKEN Research Center for Allergy and Immunology. The mouse strain, B6.129S5-*Il17rb^tm1Lex^*/Mmucd (RRID:MMRRC_032383-UCD), was obtained from the Mutant Mouse Resource and Research Center (MMRRC) at University of California at Davis, an NIH-funded strain repository, and was donated to the MMRRC by Genentech, Inc. All animals were maintained in the vivarium of the Bioscience Research Building at the University of Texas at El Paso on a 12 h light/dark cycle with ad libitum access to food and water.

### Bacteria

*Francisella tularensis* subsp. *holarctica* LVS (BEI Resources) cultures were initiated by inoculating bacterial suspensions onto Mueller–Hinton agar plates supplemented with 1% IsovitaleX (BD Bioscience) and 5% defibrinated sheep blood (Quad Five) to support optimal growth. Plates were incubated for 48 h at 37°C under standard aerobic conditions. Post-incubation, bacterial colonies were harvested, aliquoted, and diluted as required. For long-term preservation, bacterial suspensions were prepared in a cryoprotectant solution consisting of 15% glycerol in phosphate-buffered saline (PBS). Aliquots were stored at −80°C to maintain viability and stability until use. Bacterial concentration (CFU) was determined from 4 randomly selected aliquots by serial dilution plating every 9 months.

### LVS Challenge

50 µL containing 1 × 10^6^ CFU LVS diluted in sterile PBS was injected intradermally into the flank above the hindquarters of 8-12 week old mice anesthetized with 3–5% isoflurane inhalation. Animals were monitored every 12 h post-infection for the first 48 h and every 8 h thereafter. All animals were weighed before inoculation and every morning thereafter. An animal was considered terminal and humanely euthanized per AVMA standards when it had lost 20% of its baseline weight. In addition, animals were checked for clinical symptoms of disease and considered terminal when lethargic and immobile with prodding. Significance was calculated by Mantel-Cox test. Blood was drawn on days 0, 1, 3, 5, and 7 after infection from the retro-orbital capillary sinus using heparinized capillary tubes, and at the time of euthanasia (T) by cardiac puncture. Since each animal was bled every other day, alternating eyes were used to prevent irritation and ocular damage. Whole blood was fractionated and plasma frozen until completion of the experiment for subsequent analyses.

### Flow cytometry

Spleens were processed using a glass Dounce homogenizer or spleen screen. Lungs were processed using the Mouse Lung Dissociation Kit and GentleMACS Dissociator (Miltenyi Biotec: 130-093-235) according to the manufacturer’s protocol. Red blood cells were lysed using Ammonium-Chloride-Potassium (ACK) lysis buffer for 30 s, followed by inactivation with FACS buffer (PBS + 4% FBS) containing 10 µg/mL brefeldin A. Cell suspensions were filtered through a 70 µm cell strainer to remove debris. Single cell suspensions were incubated with Fc block/Fc Shield (Tonbo Bioscience: 70-0161-U100) for 15 min at 4°C to prevent nonspecific antibody binding. Cells were washed and incubated with αGalCer-loaded CD1d dextramer (Immudex: YD08002) for 10 min at 25°C to label NKT cell populations. Following dextramer labeling, cells were stained with fluorescently conjugated antibodies targeting surface markers for 20 min at 4°C. For intracellular cytokine analysis, cells were stained using Cytofix/Cytoperm fixation and permeabilization buffers (BD Biosciences: 554714) to detect IFN-γ and TNF-α. Nuclear transcription factor staining was performed using the True-Nuclear Transcription Factor Buffer Set (BioLegend: 424401) according to the manufacturer’s instructions. Samples were analyzed using a Beckman Coulter Gallios flow cytometer, and data were processed using FlowJo 10.1 (Waters Biosciences). Because no single positive marker identifies all type II NKT cells, we operationally defined type I NKT cells as B220− CD3+ NK1.1+ αGalCer-CD1d dextramer+ cells and type II NKT cells as B220− CD3+ NK1.1+ αGalCer-CD1d dextramer− cells, and interpreted these data in conjunction with Jα18^-/-^ and IL-17Rβ^-/-^ enrichment models.

### Multiplex cytokine quantification

Cytokine levels of LVS-infected mice or macrophages were assessed using the MilliporeSigma Mouse Th17 Milliplex 96-well assay (MT17MAG47K-PX25) according to the manufacturer’s protocol. This multiplex assay quantitatively measured IL-17E/IL-25, GM-CSF, IFN-γ, MIP-3α/CCL20, IL-1β, IL-2, IL-4, IL-5, IL-6, IL-21, IL-22, IL-28B, IL-10, IL-23, IL-12p70, IL-27, IL-13, IL-15, IL-17A, IL-17F, IL-33, IL-31, TNF-β, TNF-α, and CD40L. Significance was determined by two-way ANOVA with Bonferroni post-test at p-value < 0.05.

### Bulk NKT cell isolation

Erythrocytes were lysed from splenocyte suspensions using ACK/RBC lysis buffer for 30 s to selectively deplete red blood cells. The lysis reaction was terminated by neutralization with MACS buffer supplemented with 4% FBS and 2 mM EDTA. Cell suspensions were passed through a 70 µm cell strainer to remove debris and aggregated cells, followed by centrifugation at 300 × g for 10 min at 4°C to pellet cells. Bulk NKT cells were isolated using the Mouse NK1.1⁺ invariant NKT cell isolation kit (Miltenyi Biotech: 130-096-513) following the manufacturer’s instructions. Briefly, this consisted of two magnetic separation steps: 1) depletion of non-target cells and 2) positive selection of NK1.1⁺ cells. This purification strategy yielded a population containing both type I and type II NKT cells, collectively referred to as bulk NKT cells in subsequent analyses.

### NKT co-culture assay

Immortalized C57BL/6 bone marrow–derived macrophages (BEI Resources, NIAID) were plated in a 96-well plate at a concentration of 1-2 × 10^5^ cells per well in 200 µL DMEM containing 10% FBS and 0.1% ciprofloxacin and allowed to attach for 18 h. Macrophages were infected with *F tularensis* LVS at a multiplicity of infection (MOI) of 30 for 2 h in antibiotic-free DMEM. Following a PBS wash, infected cells were incubated for 1.5 h in DMEM containing 100 µg/mL gentamicin to kill extracellular bacteria. Infected cells were then incubated overnight in DMEM containing 10 µg/mL gentamicin to maintain intracellular infection and prevent bacterial overgrowth. NKT cells were added at various doses and treatments and incubated at 37°C with 5% CO_2_ overnight. Depleted non-target cells were added separately to infected cells at the same ratios to control for the addition of more cells. Supernatants were harvested after 24 hours and cytokines were quantified by single or multiplex ELISA. Percent inhibition was calculated as (control - experimental) / (control) * 100. Significance was determined by two-way ANOVA with Bonferroni post-test or one-way t-test, as appropriate, with p-value <0.05.

### Measurement of cell death

Macrophages were plated at a concentration of 1 × 10^5^ cells/well in a 96-well optical-bottom plate and infected at an MOI of 30. At 22 h post-infection, cells were stained with DiO (3,3B-dioctadecyloxacarbocyanine perchlorate; Invitrogen: D1125) for 2 h at 37°C protected from light. After 1.5 hours, cells were stained with propidium iodide (Invitrogen: P1304MP) for 30 min at 37°C protected from light. Plates were analyzed using a Molecular Devices ImageXpress Pico Automated Cell Imaging System to quantify the percentage of dead cells per well across conditions. Dead macrophages were identified as those exhibiting both green fluorescence and propidium iodide uptake.

### MACS-based Type I and Type II NKT cell isolation

Type I and type II NKT cells were separated through a modification of the Mouse NK1.1⁺ invariant NKT cell isolation kit (Miltenyi Biotech: 130-096-513). Splenocytes isolated from C57BL/6 mice were subjected to RBC lysis using ACK/RBC lysis buffer for 30 s, followed by neutralization with MACS buffer containing 4% FBS and 2 mM EDTA. Cell suspensions were filtered through a 70 µm cell strainer. Non-target cells were depleted with the antibody cocktail per the manufacturer’s instructions. Flow-through cells were collected and incubated with PE-conjugated αGalCer-loaded CD1d dextramer (Immudex: YD08002) and PIP phosphatase inhibitor (1:1000) for 40 min at 4°C with gentle agitation. Labeled cells were washed and incubated with anti-PE microbeads for 30 min at 4°C then passed through a positive selection MS column. The column was removed from the magnetic field and MACS buffer was used to flush and collect purified type I NKT cells. The remaining cells continued through the Mouse NK1.1⁺ invariant NKT cell isolation kit protocol to isolate NK1.1⁺ type II NKT cells. Purity was determined by flow cytometry. Supernatants were harvested after 24 hours and cytokines were quantified by single or multiplex ELISA.

### Cell-free supernatant transfer assays

Macrophages were seeded and infected as described above. After infection, 100 µL of supernatants collected from prior NKT subtype co-culture experiments were transferred to each well of the infected macrophage cultures. After a 24 h incubation at 37°C, supernatants were harvested and IL-6 concentrations were quantified by ELISA.

### Chemical fixation of NKT cells

Sorted NKT subtypes were resuspended in 100 µL of 0.15% glutaraldehyde and incubated for 30 s at room temperature. 100 µL of 0.2 M lysine solution was added to neutralize the glutaraldehyde. Cells were washed twice with DMEM to remove residual lysine and glutaraldehyde, resuspended in DMEM containing 10 µg/mL gentamicin, and co-cultured with infected macrophages under standard experimental conditions.

### Antibody blocking of cell signaling molecules

Macrophages were seeded and infected as described above. After infection and prior to the addition of NKT cells, various antibodies were administered at 1 µg/well. After a 24 h incubation at 37°C, supernatants were harvested and IL-6 concentrations were quantified by ELISA.

### Recombinant IL-22 treatment

Macrophages were seeded and infected as described above. After infection, recombinant mouse IL-22 protein (PeproTech) was added in the absence of NKT cell subsets. After a 24 h incubation at 37°C, supernatants were harvested and IL-6 concentrations were quantified by ELISA.

### ASO Silencing Treatment

Macrophages were seeded as described above and treated with 10 µM self-delivering antisense oligonucleotides (ASO) (AUM Biotech) overnight. Prior to addition of NKT cells, macrophages were washed and infected as described above. After a 24 h incubation at 37°C, supernatants were harvested and IL-6 and IL-22 concentrations were quantified by ELISA. To verify loss of macrophage IL-22 production, treated and untreated macrophages were stimulated with LPS.

### Real-time RT-PCR

RNA was extracted using a RNeasy kit (Qiagen) and analyzed by real-time RT-PCR using the CYBRFast one-step RT-qPCR kit (Tonbo Biosciences, San Diego, CA) and the StepOne Real-Time PCR System (Applied Biosystems, Foster City, CA). ΔCt values were calculated by comparison with GAPDH/actin expression levels, and ΔΔCt values were calculated by comparison with the average Ct value of uninfected macrophages; results are reported as fold changes in expression.

### Statistical analysis

Survival curves of LVS challenge mice were compared using Mantel-Cox test with 10-15 animals per group and were repeated 2-3 times. Plasma cytokine levels were analyzed either by ANOVA with Tukey’s post-test or multiparametric t test for repeated sampling measures. Levels of *in vitro* inflammation were compared by Mann-Whitney U test or ANOVA with Bonferroni post-test. For all tests, significance was determined at the p≤0.05 level. Statistical analyses and graphs were generated using GraphPad Prism.

## Results

### Intradermal LVS infection reveals a protective role for NKT cells

*Francisella tularensis* infection triggers an overwhelming release of proinflammatory and regulatory cytokines into the bloodstream making this infection a model of sepsis, CRS, and SIRS.[1], [2], [27] As such, lethality is directly associated with serum cytokine levels and various regulatory mechanisms have been sought using this infection to control the cytokine storm. NKT cells can produce inflammatory and regulatory effector mechanisms so they may contribute to or protect from hyperinflammation. In a previous study, Jα18-deficient mice had poorer survival compared with wildtype controls while CD1d-deficient mice had increased survival following intranasal instillation.[59] This was attributed to the production of IFN-γ from NKT cells in the lung. However, intradermal infection of either Jα18- or CD1d-deficient mice with a LD_50_ LVS for wildtype mice resulted in significantly poorer survival (Figure 1A). This was accompanied by significant increases in proinflammatory serum cytokine levels (IL-6, IFN-γ, and TNF-α) in mice deficient of NKT cell subsets, even three days after infection, compared with those possessing NKT cells (Figure 1B).

**Figure 1:**
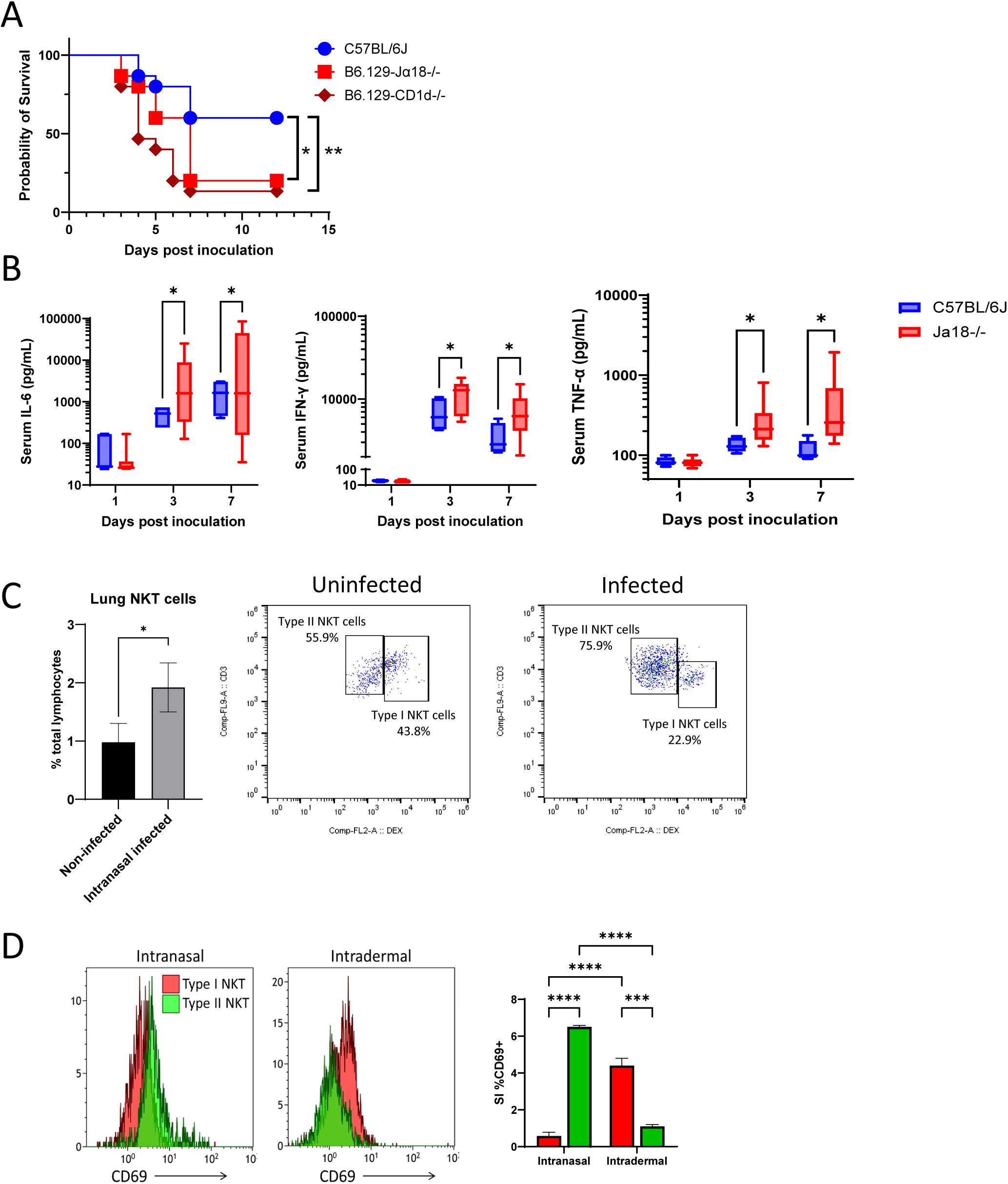
Infection route influences NKT cell subset activation favoring type II NKT cells in the lung after intranasal infection and type I NKT cells in the spleen after intradermal infection. (A) Survival of mice infected with F. tularensis LVS is significantly reduced in the absence of type I NKT cells (Jα18⁻/⁻) and total NKT cells (CD1d⁻/⁻) compared with wildtype C57BL/6J controls, *p-value <0.05, **p-value <0.01 by Mantel-Cox survival analysis. (B) Serum cytokine levels of IL-6, IFN-γ, and TNF-α are significantly suppressed by NKT cells, *p-value < 0.05 by mixed-effects analysis with Holm-Sidak multiple comparison test. (C) Intranasal instillation LVS results in increased total NKT-cell numbers and expansion of the type II NKT cell subset in the lungs of wildtype mice at 36 hours post infection, *p-value <0.05 by Mann-Whitney t test. (D) CD69 expression is likewise dependent on infection route, ***p-value <0.005 ****p-value < 0.0001 by ANOVA.

### Infection route shapes NKT-cell subset activation

To reconcile this data with prior reports, wildtype mice were infected intranasally and the lungs harvested for characterization by flow cytometry. 36 hours after instillation, we observed a significant increase in the absolute numbers of NKT cells in the lungs (Figure 1C). Subtyping revealed a profound increase in type II NKT cells (CD3+ NK1.1+ αGalCer-CD1d dextramer−) and a compensatory decrease in type I NKT cells (CD3+ NK1.1+ αGalCer-CD1d dextramer+). These type II NKT cells displayed a higher expression of CD69 compared with type I NKT cells indicating a predominant activation of type II NKT cells in the lung following intranasal LVS infection (Figure 1D). In contrast, intradermal infection with LVS resulted in greater CD69 expression on type I NKT cells.

### Type I NKT cells suppress macrophage cytokine production

The increased serum cytokine levels and worsened survival of mice deficient in NKT cells suggest that NKT cells suppress inflammation following intradermal inoculation with LVS. To test this, NKT cells were purified from the spleens of naïve mice and cultured with LVS-infected macrophages. Purification increased the population of NKT cells from <5% to ∼80% of recovered cells while non-targeted cells contained nearly no NKT cells (Figure 2A). Of note, the purified splenic NKT cell population was dominated (∼60%) by type I NKT cells. Addition of NKT cells to LVS-infected macrophages significantly reduced the production of IL-1β and IL-6, the initiator and effector cytokines of the inflammatory response, by 85-95% with no alternation of cell viability (Figure 2B, S1). Importantly, addition of an equal number of non-target cells depleted of NKT cells did not inhibit but slightly enhanced the production of these cytokines making NKT cells responsible for the suppression. Indeed, NKT cells inhibited nearly an identical percentage of cytokine compared either to that produced by infected macrophages alone or infected macrophages to which a matching number of non-target cells were cultured (Figure 2C). Multiplex analysis of culture supernatant revealed that NKT cell inhibition of inflammation was not limited to IL-1β and IL-6 but ten additional cytokines demonstrating their capacity to suppress the cytokine storm (Figure 2D).

**Figure 2:**
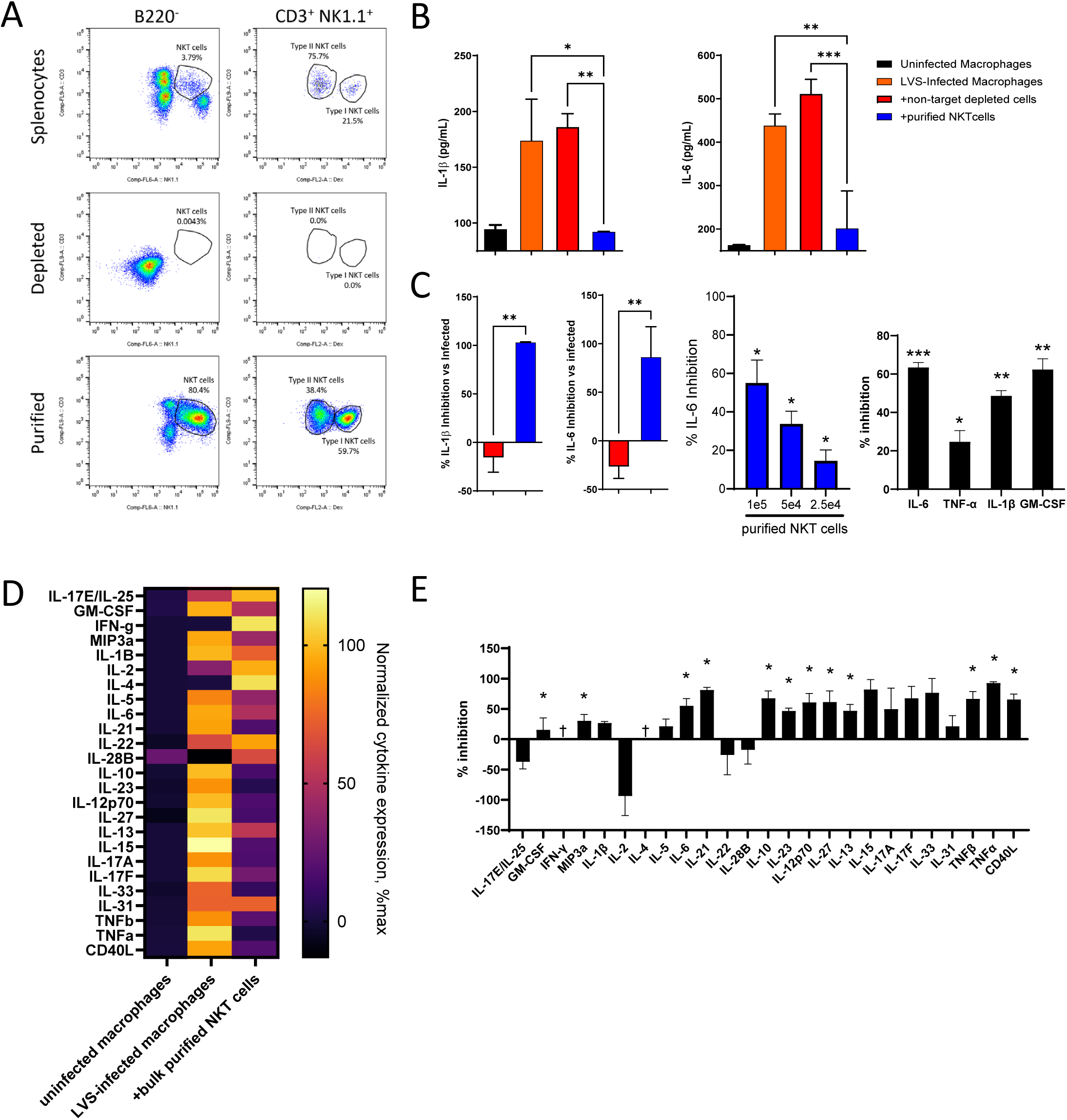
Purified total NKT cells suppress LVS-induced inflammatory cytokine production in vitro. (A) Flow cytometry analysis of whole splenocytes, NKT-depleted splenocytes, and purified NKT cells. (B-C) Co-culture of purified NKT cells with LVS-infected macrophages resulted in reduced production of IL-1β, IL-6, TNF-α, and GM-CSF whereas NKT-depleted splenocytes did not reduce these inflammatory cytokines. *p-value <0.05 **p-value < 0.01 ***p-value <0.005 by ANOVA (D, E) Multiplex analysis demonstrates that the addition of purified NKT cells to LVS-infected macrophages decreases a broad array of inflammatory and regulatory cytokines associated with the cytokine storm. (D) Heat map normalized to the minimal and maximal values for each cytokine. (E) Percent inhibition of each cytokine calculated against LVS-infected macrophages, *p-value <0.05 by one sample t test.

As type I and type II NKT cells showed differential activation following intranasal and intradermal LVS infection leading to differing outcomes, these subsets were separately purified from naïve mice and cultured with LVS-infected macrophages. Type I NKT cells inhibited the production of IL-6 in a dose-dependent manner while type II NKT cells had severely limited, if any, ability to inhibit IL-6 production (Figure 3A, S2). To further confirm that type I NKT cells suppress LVS-induced IL-6, NKT cells were purified from wildtype, Jα18-deficient and IL-17Rβ-deficient mice. These transgenic mice are deficient in type I NKT cells or severely reduced type II NKT cells, respectively. Purification of NKT cells from wildtype mice produced a mixed population of NKT cell subsets, while Jα18-deficient mice produced nearly pure type II NKT cells, and IL-17Rβ-deficient mice were dominated by type I NKT cells (Figure 3B). Cultured with LVS-infected macrophages, NKT cells purified from wildtype animals and IL-17Rβ-deficient animals, both populations of which had high proportions of type I NKT cells, were able to significantly inhibit LVS-induced IL-6 while the type II NKT cell-dominated population from Jα18-deficient mice could not inhibit IL-6 production. Like the bulk purified NKT cells (Figure 2D), purified type I NKT cells inhibited an additional nine cytokines found in culture supernatants, including numerous pro-inflammatory cytokines (Figure 3C, S3).

**Figure 3:**
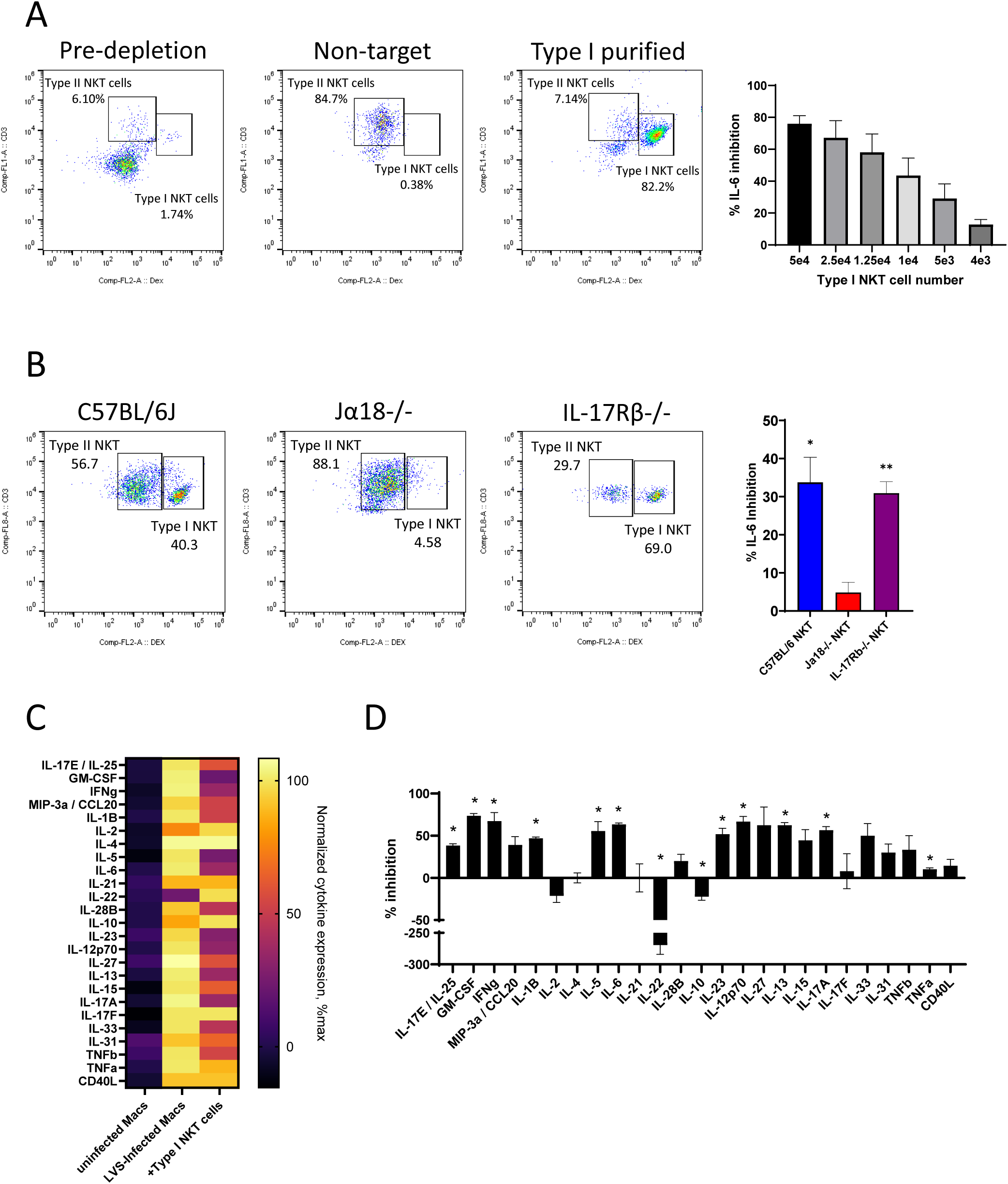
Type I NKT cells inhibit cytokine storm cytokine production. (A) Type I NKT cells purified from wildtype mice inhibit IL-6 production by LVS-infected macrophages in a dose-dependent manner. (B) NKT cell populations purified from transgenic mice deficient in either type I (Jα18⁻/⁻) or type II (IL-17Rβ⁻/⁻) NKT cells inhibit IL-6 production from LVS-infected macrophages only when type I NKT cells are present, **p-value <0.005 ***p-value <0.001 by ANOVA with Bonferroni posthoc test. (C, D) Multiplex analysis of type I NKT cells co-cultured with LVS-infected macrophages demonstrates inhibition across a broad range of proinflammatory and regulatory cytokines. (D) Heat map normalized to the minimal and maximal values for each cytokine. (E) Percent inhibition of each cytokine calculated against LVS-infected macrophages, *p-value <0.05 by one sample t test.

### Type I NKT cell suppression is mediated by soluble IL-22

Glutaraldehyde fixation of these inhibitory type I NKT cells prior to their addition to LVS-infected macrophages eliminated their ability to inhibit inflammation, suggesting a secreted effector molecule or dynamic cell surface protein (Figure 4A). Transfer of cell-free conditioned supernatant to fresh LVS-infected macrophages transferred the ability of type I NKT cells to inhibit inflammation, demonstrating the presence of a soluble effector molecule. Transfer of similar conditioned supernatant from control cells did not inhibit IL-6 production. Multiplex analysis of the culture supernatant of inhibitory type I NKT cells and fixed NKT cells revealed the production of IL-22 only when NKT cells were added to the culture, the production of which was prevented by prior fixation of type I NKT cells (Figure 3C, S3). The production and loss of IL-22 in purified versus fixed NKT cells directly correlated with NKT cell inhibitory activity (Figure S4A). Neutralization of a variety of molecular targets did not affect IL-6 suppression, except for IL-22 and IL-1β (which is directly required for the production of IL-6) (Figure S4B). Addition of αIL-22 prevented the accumulation of IL-22 in the culture supernatant resulting in no inhibition by type I NKT cells (Figure 4C). In contrast, addition of recombinant IL-22 at concentrations observed in type I NKT cell cultures was sufficient to inhibit IL-6 production in NKT cell-free LVS-infected macrophage cultures.

**Figure 4:**
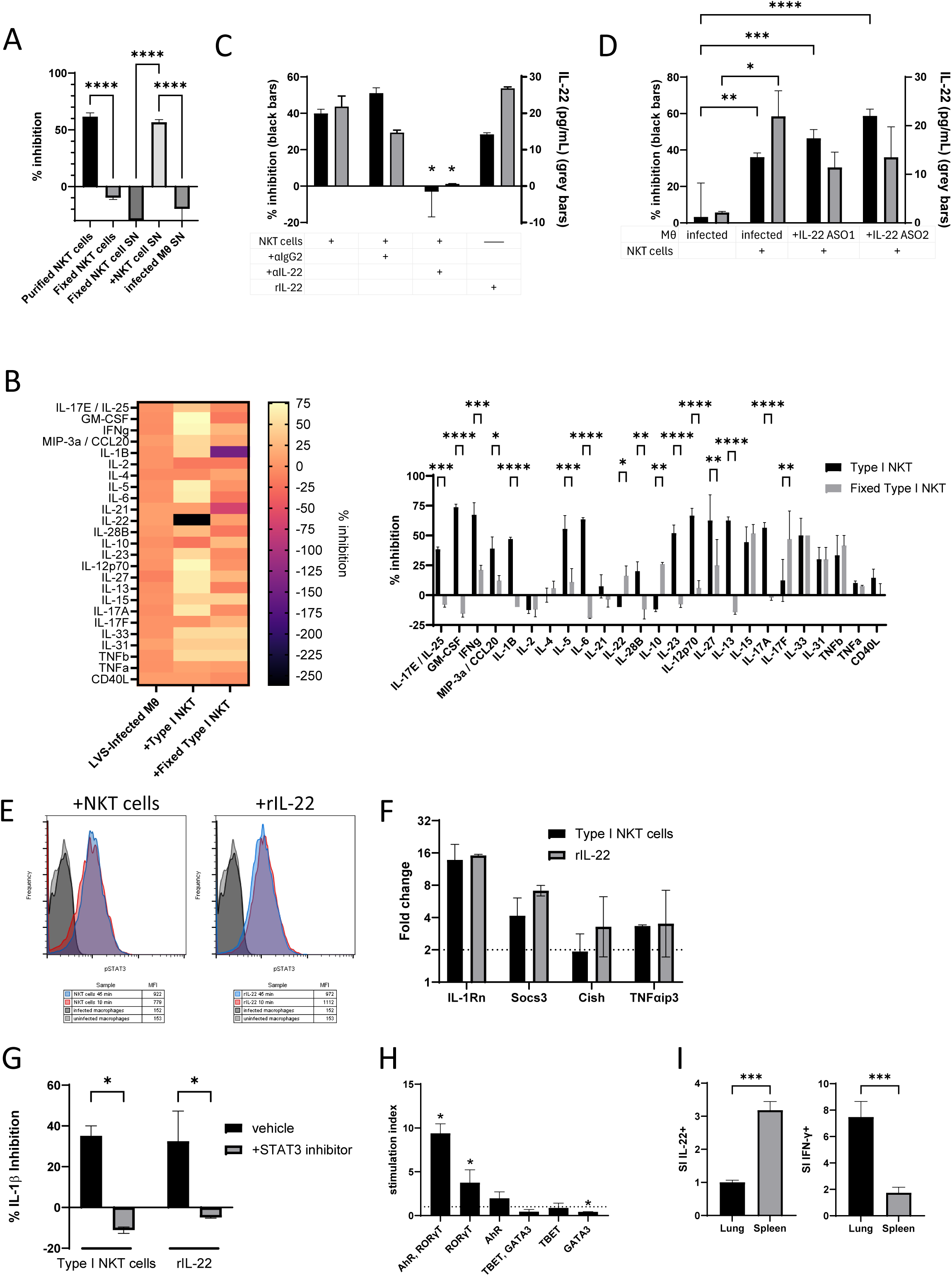
Type I NKT-cell–derived IL-22 inhibits cytokine-storm–associated cytokine production. (A) Glutaraldehyde-fixed type I NKT cells cannot inhibit IL-6 production from LVS-infected macrophages. Supernatants from previously activated NKT cells alone can inhibit IL-6 production in NKT cell-free cultures whereas supernatants from fixed type I NKT cells fails to inhibit IL-6 production. (B) Multiplex cytokine profiling revealed inhibition of multiple cytokine storm-associated cytokines, except for IL-22. IL-22 production is directly associated with functional type I NKT cell-mediated cytokine inhibition. (C) Anti-IL-22 neutralization depletes IL-22 from the supernatant (grey bars), eliminating IL-6 inhibition (black bars) by type I NKT cells. (D) IL-22 antisense oligonucleotide (ASO) silencing in macrophages prior to NKT cell co-culture does not disrupt IL-6 inhibition (black bars) or IL-22 levels (grey bars). (E) Both type I NKT cells and recombinant IL-22 stimulate phosphorylation of STAT3 in LVS-infected macrophages within 10–45 minutes as detected by flow cytometry. **p-value <0.005 ***p-value <0.001 ****p-value <0.0001 by ANOVA. (F) Type I NKT cells induce transcription of STAT3-regulated genes (IL-1Rα, Socs3, Cish, Tnfaip3) in LVS-infected macrophages within 2.5 hours similar to addition of recombinant IL-22. (G) Pharmacologic inhibition of STAT3 restores IL-1β produced within 2.5 hours from LVS-infected macrophages cultured with type I NKT cells or recombinant IL-22. *p-value <0.05 by ANOVA. (H) Flow cytometric quantification of transcription factors expressed by spleen-derived NKT cells 36 hours after LVS infection of C57BL/6 mice shows expansion of AhR- and RORγT-positive NKT cell populations. *p-value <0.05 by one sample t test. (I) Intradermal LVS infection stimulates IL-22 expression in splenic NKT cells but not in lung NKT cells which instead express IFN-γ. ***p-value <0.001 by ANOVA.

Because IL-22 was detected in co-culture and macrophages could produce IL-22 after LPS stimulation (Figure S4C), we tested whether macrophage-derived IL-22 accounted for IL-6 suppression. Silencing of IL-22 mRNA translation using self-delivering antisense oligonucleotides (ASO) reduced macrophage IL-22 production in the LPS control but did not reduce IL-22 accumulation or IL-6 suppression when ASO-treated macrophages were cultured with type I NKT cell (Figure 4D). Conversely, ASO silencing of IL-22 in type I NKT cells prior to co-culture proved toxic to the NKT cells. Taken together, these data support type I NKT cells as the dominant IL-22 source under these conditions.

### IL-22 activates STAT3-dependent anti-inflammatory programs in macrophages

IL-22 binding initiates JAK1/Tyk2-mediated signal transduction through phosphorylation of STAT3 in macrophages [71], [72]. Type I NKT cells triggered phosphorylation of STAT3 apparent 10-45 minutes after addition, similar to that observed following addition of rIL-22 (Figure 4E). Activated pSTAT3 initiated transcription of the signature genes *IL-1Rα*, *Socs3*, *Cish,* and *TNFAip3* detectable 2.5 hours after addition of type I NKT cells similar to addition of rIL-22 (Figure 4F). Ultimately, activation of STAT3 signaling by either type I NKT cells or rIL-22 inhibited the production of IL-1β which was blocked by addition of a STAT3 inhibitor (Figure 4G). *In vivo*, LVS infection enhanced expression of the AhR and RORγT transcription factors in splenic NKT cells compared with a reduction TBET- and GATA3-expressing cells (Figure 4H) supportive of an IL-22-permissive splenic NKT-cell phenotype.

These data suggest that splenic type I NKT cells produce IL-22 to reduce levels of inflammatory cytokines produced by macrophages. Indeed, following intradermal LVS infection, there was a 3-fold increase in splenic NKT cells producing IL-22 (Figure 4I). Meanwhile, lung NKT cells instead produced IFN-γ (Figure 4I) following intranasal instillation, which correlated with the pro-inflammatory environment previously described.[59] These data demonstrate that LVS-stimulated splenic type I NKT cells act to control inflammation and the cytokine storm through an IL-22-mediated effector mechanism.

## Discussion

Cytokine storm-driven pathology remains a central determinant of morbidity and mortality during severe infection, sepsis, and related inflammatory syndromes. Although NKT cells respond rapidly during infection, their precise role in either amplifying or restraining hyperinflammation, whether sterile or responsive to infection, has remained controversial. Prior studies have frequently treated NKT cells as a single functional population. The present study demonstrates that type I (invariant) NKT cells function as dominant suppressors of infection-induced hypercytokinemia, whereas type II NKT cells provide minimal protection in the context of intradermal *Francisella tularensis* infection. Importantly, we identify IL-22 as the key soluble effector molecule mediating this immunosuppressive function, thereby defining a previously unrecognized type I NKT cell–IL-22–STAT3 axis that restrains pathological inflammation.

Our in vivo data demonstrate that loss of NKT cells, either globally (CD1d deficiency) or selectively (Jα18 deficiency), results in worsened survival and sustained elevation of systemic pro-inflammatory cytokines following intradermal LVS challenge. These findings do not contradict prior intranasal infection studies, but instead suggest that NKT cell function is shaped by anatomic site and route of infection. Intranasal LVS favors pulmonary type II NKT cell activation and IFN-γ production, whereas intradermal LVS favors splenic type I NKT cell activation and IL-22-associated suppression ofsystemic cytokines.

These observations align with emerging evidence that type I and type II NKT cells frequently exert opposing immunological roles, shaped by tissue localization, antigen availability, and cytokine milieu rather than the structure of their receptor. Type I NKT cells have been shown to promote immune regulation in models of autoimmunity and tissue injury, while type II NKT cells can either suppress or exacerbate inflammation depending on context. Our data extend this paradigm to cytokine storm biology and demonstrate that failure to separate NKT cell subsets can obscure fundamentally divergent functions.

Using an in vitro co-culture system, we show that purified NKT cells potently suppress production of IL-1β, IL-6, TNF-α, and additional inflammatory mediators from LVS-infected macrophages. This effect was dose-dependent, transferable via conditioned supernatant, and abolished by chemical fixation of type I NKT cells. Collectively, these data indicate suppression by secreted effector molecules rather than cytotoxicity or contact-dependent inhibition. Importantly, this suppressive capacity was confined almost entirely to type I NKT cells as purified type II NKT cells failed to significantly inhibit IL-6 production.

These findings challenge the traditional view that early innate-like lymphocyte responses predominantly amplify inflammation and instead position type I NKT cells as critical negative regulators of the macrophage-driven cytokine cascades during severe bacterial infection. Given the central role ofmacrophages as initiators and amplifiers ofcytokine storm, this regulatory interaction likely represents a key checkpoint limiting progression from protective inflammation to systemic pathology.

These data identify IL-22 as a necessary and sufficient mediator of type I NKT cell suppression of macrophage IL-6 and IL-1β in this in vitro co-culture system and suggest that IL-22 contributes to the cytokine suppression induced by type I NKT cells. Neutralization of IL-22 completely abrogated NKT-dependent inhibition of IL-6, while addition of recombinant IL-22 at concentrations produced by NKT cells was sufficient to recapitulate this effect in the absence of NKT cells. Moreover, type I NKT cell–derived IL-22 activated canonical STAT3 signaling in macrophages, inducing transcription of well-established anti-inflammatory and cytoprotective genes including il1rn, socs3, cish, and tnfaip3.

IL-22 is traditionally associated with epithelial barrier protection, tissue repair, and antimicrobial defense at mucosal surfaces. However, increasing evidence suggests that IL-22 also exerts direct immunomodulatory effects on myeloid cells, particularly through STAT3-dependent negative feedback mechanisms. Our findings place type I NKT cells among the physiologically relevant cellular sources of IL-22 during systemic infection and highlight a previously unrecognized role for IL-22 in actively suppressing macrophage cytokine storm responses, rather than merely promoting tissue repair.

Notably, IL-22 production by NKT cells was restricted to splenic but not pulmonary compartments, correlating with reduced inflammation systemically but heightened inflammation in the lung. This tissue tropism was associated with increased RORγT expression and reduced TBET and GATA3 expression in splenic NKT cells, consistent with a transcriptional program permissive for IL-22 production. These findings reinforce the concept that NKT cell functional polarization is highly tissue-dependent and that the local milieu shapes their effector repertoire during infection.

The absence of IL-22 production by lung NKT cells may help explain why respiratory infection models often implicate NKT cells in immunopathology, whereas peripheral infections reveal protective roles. Therapeutic strategies aimed at selectively enhancing IL-22–producing type I NKT cells or mimicking their downstream signaling pathways, may therefore require careful consideration of tissue context, perhaps with directed delivery.

This study does have recognized limitations for the totality of the NKT cell-IL-22-STAT3 axis. First, type II NKT cells were operationally defined as CD3+NK1.1+αGalCer-CD1d dextramer-negative cells, a population that may include additional innate-like T cells whose role is not accounted for. Second, IL-22 dependence was defined most directly in in vitro co-culture assays; future in vivo IL-22 blockade or cell-specific IL-22 loss-of-function studies will be needed to determine the contribution of this pathway to survival and systemic cytokine control during intradermal LVS infection. Third, the use of immortalized bone marrow-derived macrophages provides a controlled reductionist system, while reducing dependence on animal research, but may not fully capture tissue macrophage heterogeneity during infection.

In summary, this study identifies type I NKT cells as suppressors of intradermal F. tularensis-induced hyperinflammation and defines type I NKT cell-IL-22-dependent activation of macrophage STAT3 signaling as a key mechanism limiting inflammatory cytokine production. These findings help reconcile prior reports of pathogenic pulmonary NKT cell activity by showing that NKT cell function depends on infection route, tissue compartment, and subset composition. More broadly, the data support a model in which early innate-like lymphocyte responses do not uniformly amplify inflammation but can instead impose cytokine-specific regulatory checkpoints that limit progression from protective antimicrobial immunity to systemic immunopathology.

## Author Contributions

All authors directly generated data presented in this manuscript. CMT, NRS and CTS contributed to writing and editing the manuscript. CTS obtained funding and oversaw the project scope.

## Acknowledgements

Supplemental Figure 1: Type I NKT cells do not induce cytotoxicity in LVS-infected macrophages. Cell death was visualized by DiO (3,3Ö-dioctadecyloxacarbocyanine perchlorate) macrophage staining (green) and propidium iodide (red) staining. Quantification of macrophage death is shown in the accompanying bar graph, representing the percentage of dead cells defined as DiO-positive macrophages that also exhibit propidium iodide fluorescence.

Supplemental Figure 2: Purified type II NKT cells exhibit minimal inhibition of IL-6 production in LVS-infected macrophages. MACS-purified type II NKT cells were added to LVS-infected macrophage cultures and assessed for their ability to inhibit IL-6 production. In contrast to type I NKT cells, type II NKT cells demonstrated only minimal inhibitory activity.

Supplemental Figure 3: Type I NKT cells reshape the cytokine environment during LVS infection, dampening proinflammatory outputs while driving IL-22 production. Line graph connecting nodes from multiplex cytokine analysis. Infection with LVS drove cytokine expression, the majority of which were inhibited by addition of type I NKT cells. IL-22 production is highlighted in red showing an increase only when functional NKT cells are added.

Supplemental Figure 4. Type I NKT cell-mediated inhibition of inflammatory cytokines is tied to their capacity to produce IL-22. (A) Suppression of IL-6 by type I NKT cells is directly correlated with IL-22 production. (B) Neutralizing antibody blockade of multiple cytokines and receptors produced no loss of IL-6 suppression, except for IL-22 and IL-1β. (C) Antisense oligonucleotides (ASO) disrupt IL-22 production from macrophages stimulated with LPS.

## Notes

### Competing Interest Statement

The authors have declared no competing interest.

### Summary of Updates

Textual revisions for clarity and figure legends.

### Funding

- National Institute of Allergy and Infectious Diseases: 1SC1AI148753

