## Supplementary figures and images for "Type I Natural Killer T Cells Suppress Infection-Induced Cytokine Storm Through an IL-22-STAT3 Axis"

### Supplementary figures 1-4

#### Supplemental Figure S1

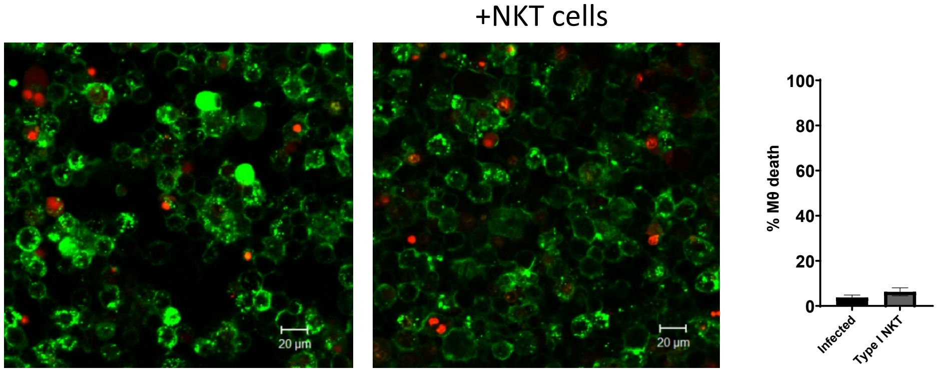

#### Supplemental Figure S2

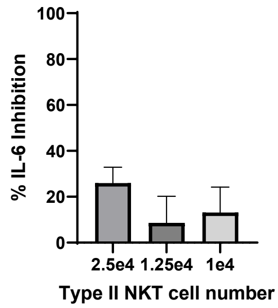

#### Supplemental Figure S3

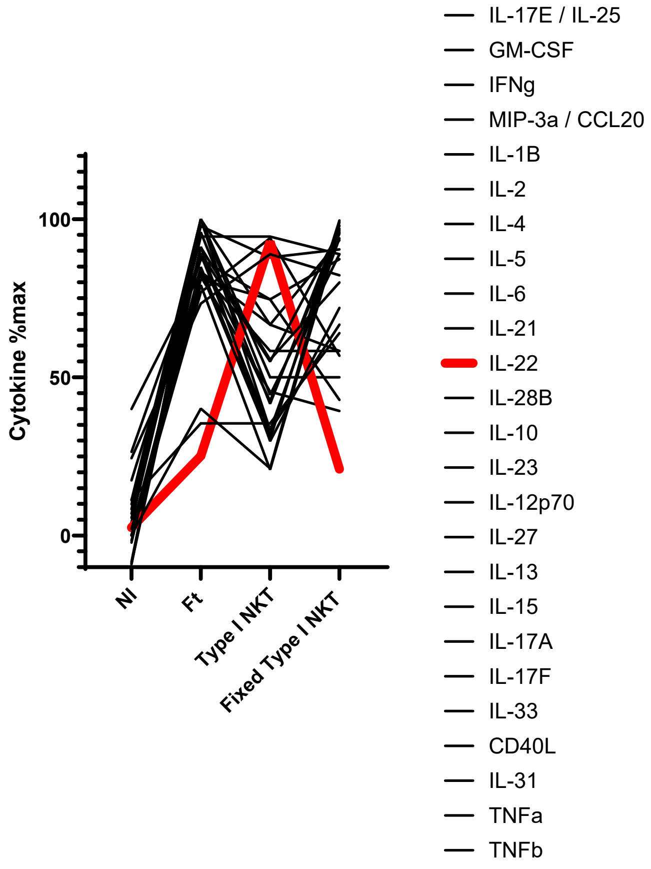

#### Supplemental Figure S4

A

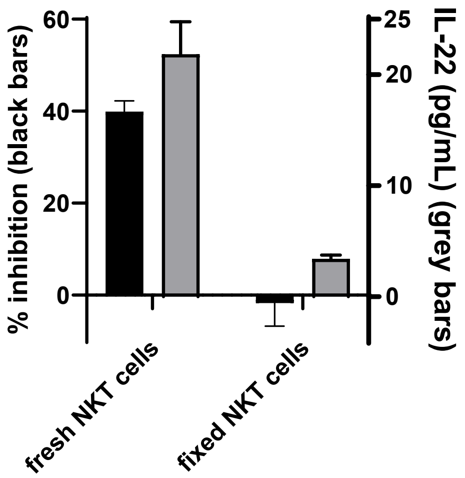


C

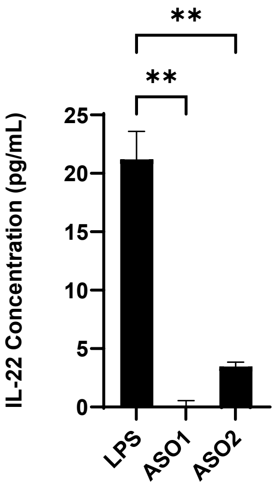


B

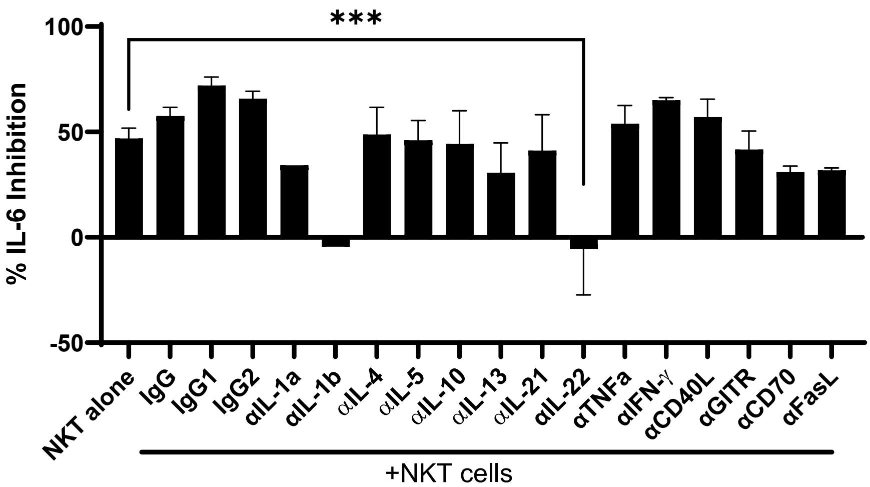
